# Dbx1 pre-Bötzinger complex interneurons comprise the core inspiratory oscillator for breathing in adult mice

**DOI:** 10.1101/271585

**Authors:** Nikolas C. Vann, Francis D. Pham, Kaitlyn E. Dorst, Christopher A. Del Negro

## Abstract

The brainstem pre-Bötzinger complex (preBötC) generates inspiratory breathing rhythms, but which neurons comprise its rhythmogenic core? Dbx1-derived neurons may play the preeminent role in rhythm generation, an idea well founded at perinatal stages of development but not in adulthood. We expressed archaerhodopsin or channelrhodopsin in Dbx1 preBötC neurons in intact adult mice to interrogate their function. Prolonged photoinhibition slowed down or stopped breathing, whereas prolonged photostimulation sped up breathing. Brief inspiratory-phase photoinhibition evoked the next breath earlier than expected, whereas brief expiratory-phase photoinhibition delayed the subsequent breath. Conversely, brief inspiratory-phase photostimulation increased inspiratory duration and delayed the subsequent breath, whereas brief expiratory-phase photostimulation evoked the next breath earlier than expected. Because they govern the frequency and precise timing of breaths in awake adult mice with sensorimotor feedback intact, Dbx1 preBötC neurons constitute an essential core component of the inspiratory oscillator, knowledge directly relevant to human health and physiology.

## INTRODUCTION

Inspiratory breathing movements in mammals originate from neural rhythms in the brainstem preBötzinger Complex (preBötC) (Feldman et al., 2013; Smith et al., 1991). Although the preBötC has been identified in a range of mammals including bats, moles, goats, cats, rabbits, rats, mice, and humans (Mutolo et al., 2002; Pantaleo et al., 2011; Ruangkittisakul et al., 2011; Schwarzacher et al., 1995, 2010; Smith et al., 1991; Tupal et al., 2014; Wenninger et al., 2004) its neuronal constituents remain imprecise. Competing classification schemes emphasize peptide and peptide receptor expression (Gray et al., 1999, 2001; Stornetta et al., 2003a; Tan et al., 2008) as well as a glutamatergic transmitter phenotype (Funk et al., 1993; Stornetta et al., 2003b; Wallen-Mackenzie et al., 2006) as cellular markers that define the preBötC rhythmogenic core.

Interneurons derived from precursors that express the homeodomain transcription factor Dbx1 (i.e., Dbx1 neurons) also express peptides and peptide receptors associated with respiratory rhythmogenesis, and are predominantly glutamatergic. *Dbx1* knock-out mice die at birth of asphyxia and the preBötC never forms (Bouvier et al., 2010; Gray et al., 2010). In rhythmically active slice preparations from neonatal Dbx1 reporter mice, Dbx1 preBötC neurons discharge in bursts in phase with inspiration (Picardo et al., 2013), and their sequential laser ablation slows and then stops respiratory motor output (Wang et al., 2014). These results obtained from perinatal mice suggest that Dbx1 neurons comprise the rhythmogenic preBötC core, i.e., the Dbx1 core hypothesis.

Nevertheless, in addition to their putatively rhythmogenic role, Dbx1 preBötC neurons also govern motor pattern. Hypoglossal motoneurons that maintain airway patency receive rhythmic synaptic drive from Dbx1 neurons within the preBötC and adjacent intermediate reticular formation (Revill et al., 2015; Song et al., 2016; Wang et al., 2014). In anesthetized vagotomized adult mice, photostimulation of Dbx1 preBötC neurons modulates inspiratory timing and its motor pattern, which is mediated in part by somatostatin-expressing (Sst) preBötC neurons (Cui et al.,2016), a large fraction of which are derived from Dbx1-expressing progenitors (Bouvier et al., 2010; Gray et al., 2010; Koizumi et al., 2016).

In adult animals, Dbx1 preBötC neurons serve non-respiratory roles as well. A subset that expresses Cadherin-9 (Cdh9) projects to the pontine locus coeruleus to influence arousal (Yackle et al., 2017). Collectively, the fractions of motor output-related (Sst-expressing) and arousal-related (Cdh9-expressing) Dbx1 neurons could account for 73% of Dbx1 neurons within the preBötC: up to 17% of Dbx1 preBötC neurons express Sst and 56% express Cdh9 with no overlap between Sst and Cdh9 expression (Bouvier et al., 2010; Cui et al.,2016; Gray et al., 2010; Yackle et al., 2017). That accounting would leave 27% of Dbx1 preBötC neurons exclusively rhythmogenic, if one assumes that all remaining Dbx1 neurons are dedicated to respiration and that single Dbx1 preBötC neurons cannot fulfill multiple duties. Therefore, while their rhythmogenic role is well established at perinatal stages of development (Bouvier et al., 2010; Gray et al., 2010), the contemporary studies recapped above from adult mice imply that rhythm generation may not be the principal function of Dbx1 preBötC neurons.

Here we reevaluate the inspiratory rhythmogenic role of Dbx1 preBötC neurons in adult mice with intact sensorimotor feedback. Using optogenetic technologies to photoinhibit or photostimulate Dbx1 neurons, we show that their perturbation affects breathing frequency and the precise timing of individual breaths within the breathing cycle, which are key properties of a core oscillator microcircuit. Other respiratory and non-respiratory roles notwithstanding, these data indicate that Dbx1 preBötC neurons constitute an essential core oscillator for inspiration.

## RESULTS

### ArchT activation hyperpolarizes Dbx1 preBötC neurons postsynaptically

We illuminated the preBötC in transverse medullary slices from neonatal Dbx1;ArchT mice (the intersection of a *Dbx1*^CreERT2^ driver mouse and a reporter featuring Cre-dependent archaerhodopsin [ArchT] expression) that spontaneously generate inspiratory rhythm and airway-related hypoglossal (XII) motor output. Light application (589 nm) to the preBötC bilaterally stopped rhythm and motor output at all light intensities (Figure 1 – figure supplement 1A and B). Dbx1 preBötC neurons recorded in whole-cell patch-clamp hyperpolarized 6.5 ± 1.0, 8.1 ± 1.1, and 11.0 ± 2.5 mV in response to light of increasing intensity (Figure 1A, cyan). We reapplied the highest intensity light in the presence of TTX, which hyperpolarized Dbx1 preBötC neurons by 8.6 ± 1.4 mV (Figure 1A and B, cyan). Light-evoked hyperpolarization was commensurate before and after TTX (Mann-Whitney U, p = 0.2, n_1_ = 8, n_2_ = 3), which suggests that ArchT hyperpolarizes Dbx1 preBötC neurons via direct postsynaptic effects.

**Figure 1.**
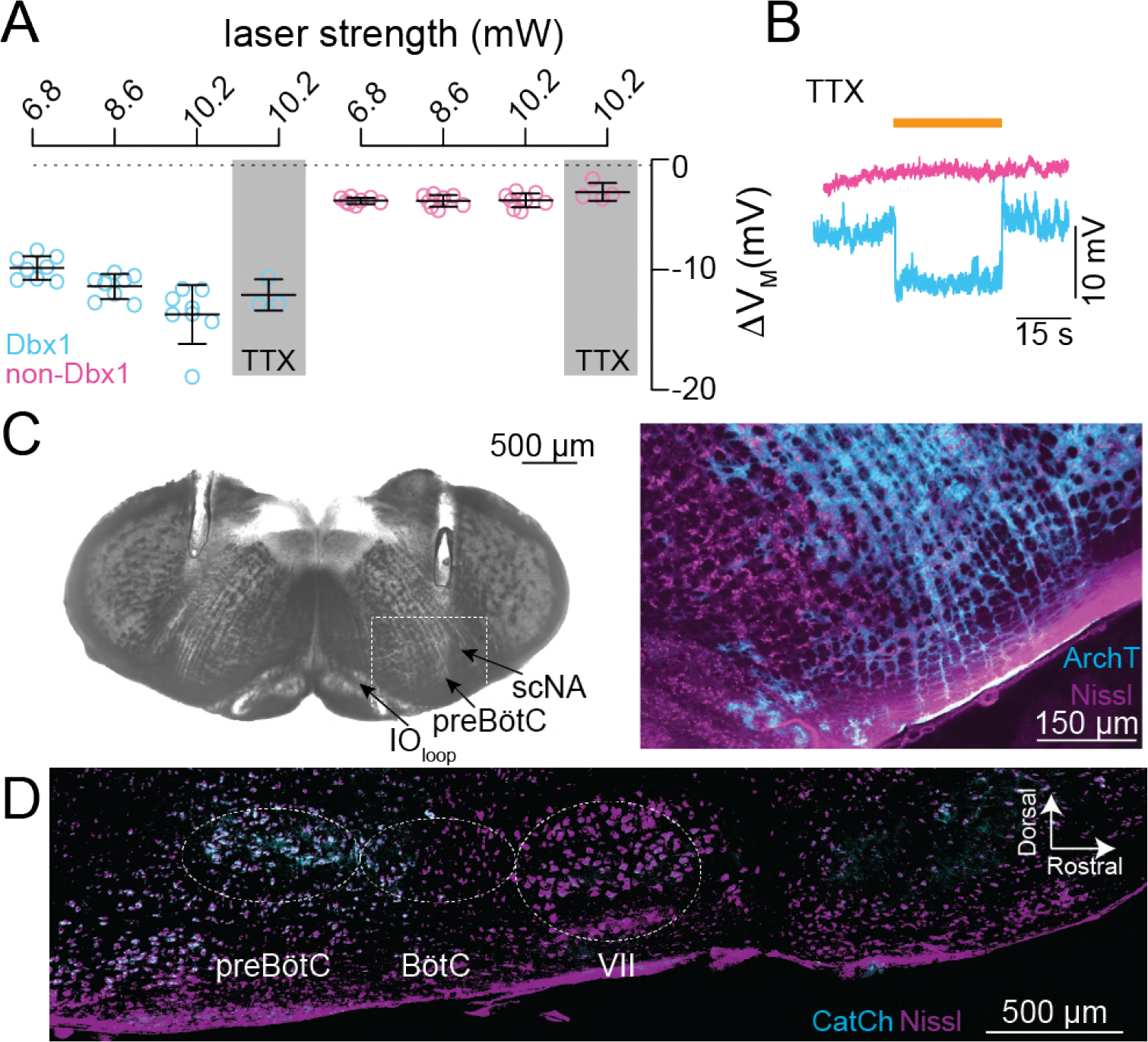
Photoinhibition of preBötC neurons *in vitro* and fusion protein expression patterns. **A**, Membrane hyperpolarization (ΔV_M_) evoked by light pulses at three intensities in Dbx1 and non-Dbx1 preBötC neurons recorded from neonatal Dbx1;ArchT mouse slices. Bars show mean and SD (n = 8 Dbx1 neurons in control and n = 4 Dbx1 neurons in 1 μM tetrodotoxin [TTX]; n = 8 non-Dbx1 neurons in control and n = 4 non-Dbx1 neurons in TTX). **B**, Membrane trajectories in response to 30-s bouts of 10.2 mW illumination in 1 μM TTX. **C**, Bright field image of a transverse section from an adult Dbx1;ArchT mouse at the level the preBötC, as indicated by the loop of the inferior olive (IO_loop_) and the semi-compact division of the nucleus ambiguus (scNA). Parallel tracks of implanted fiber optics are visible from the dorsal border of the tissue section into the intermediate reticular formation dorsal to the preBötC. The selection box was imaged using fluorescence microscopy to show ArchT (cyan) protein expression in the preBötC in detail, Nissl staining (magenta) included for contrast. **D**, Parasagittal section from an adult Dbx1;CatCh mouse. Nissl (magenta) shows anatomical landmarks including the facial (VII) cranial nucleus, Bötzinger complex (BötC), and the preBötC. CatCh (cyan) expression is limited to the preBötC.

In the same slices from neonatal Dbx1;ArchT mice, we illuminated the preBötC bilaterally while patch recording neighboring non-Dbx1 preBötC neurons. Baseline membrane potential in non-Dbx1 preBötC neurons responded negligibly to light, hyperpolarizing 0.7 ± 0.3, 1.1 ± 0.5, and 1.1 ± 0.6 mV in response to light of increasing intensity (Figure 1A, magenta, and Figure 1 – figure supplement 1B). In TTX, light at the highest intensity hyperpolarized non-Dbx1 neurons by 0.3 ± 0.8 mV (Figure 1A and B, magenta), which was indistinguishable from light-evoked hyperpolarization before TTX application (Mann-Whitney U, p = 0.2, n_1_ = 8, n_2_ = 4). These results suggest that light-evoked cessation of inspiratory rhythm and motor output *in vitro* is largely attributable to direct postsynaptic effects on Dbx1 preBötC neurons rather than network disfacilitation, which would comparably affect Dbx1 as well as non-Dbx1 neurons in the preBötC and would be eliminated by TTX.

### Photoinhibition of Dbx1 preBötC neurons attenuates breathing and resets inspiration

Next we illuminated the preBötC bilaterally using fiberoptic implants (Figure 1C shows tracks of fiberoptics in post-hoc histology) in sedated adult Dbx1;ArchT mice, which reduced breathing in all instances (Figure 2A). In control conditions breathing frequency (*f*) was typically ~3.5 Hz, tidal volume (V_T_) was ~0.1 ml, and minute ventilation (MV) was ~50 ml/min. The lowest intensity light (6.8 mW) decreased *f* by 0.3 Hz (t-test, p = 0.05, n = 6), decreased V_T_ by 0.2 ml (but that change was not statistically significant by t-test, p = 0.06, n = 6), and decreased MV by 9 ml/min (t-test, p = 0.01, n = 6) (Figure 2B).

**Figure 2.**
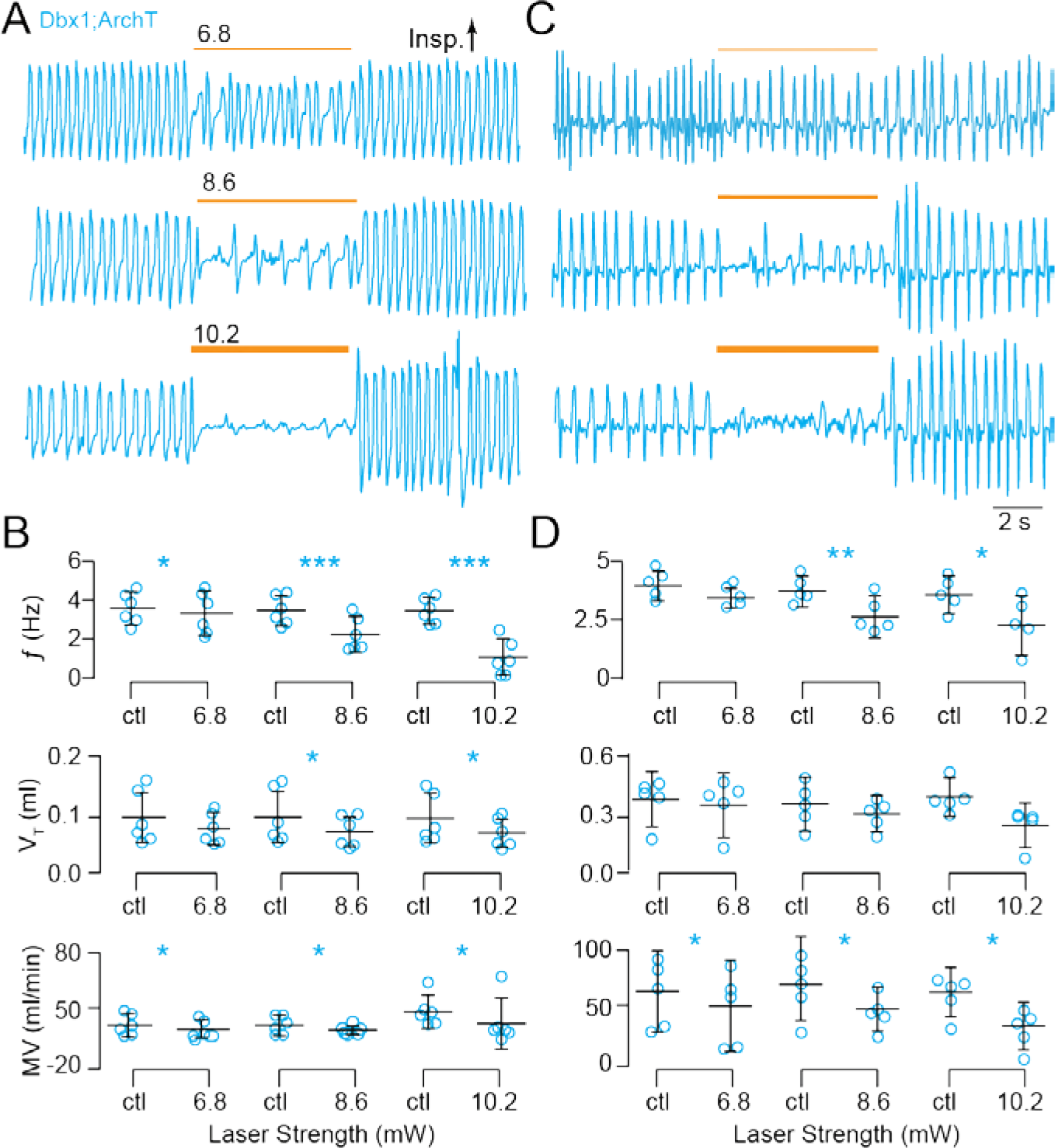
Photoinhibition of Dbx1 preBötC neurons depresses breathing in adult Dbx1;ArchT mice. **A**, Airflow traces from a sedated mouse exposed to 5-s bouts of bilateral preBötC illumination at three intensities (units of mW). Yellow line thickness corresponds to light intensity, which is also annotated above each line. **B**, Group data from experiments in A quantifying light-evoked changes in *f*, V_T_ and MV. Symbols show the mean *f*, V_T_, and MV measured in each mouse. Bars show the mean and SD for all animals tested (n = 5). Control measurements are labeled ‘ctl’; numerals indicate light intensity. **C**, Airflow traces from an awake unrestrained mouse exposed to 5-s bouts of bilateral preBötC illumination at three intensities. Yellow line thickness corresponds to light intensity; annotations mach those in A. **D**, Group data from experiments in C quantifying light-evoked changes in *f*, V_T_ and MV. Symbols show the mean *f*, V_T_, and MV measured in each mouse. Bars show the mean and SD for all animals tested (n = 6). Control measurements are labeled ‘ctl’; numerals indicate light intensity. Asterisks represent statistical significance at p < 0.05; the double asterisk represents p < 0.01; and triple asterisks represent p < 0.001.

*f*, V_T_, and MV decreased to a greater extent in response to 8.6 and 10.2 mW intensity illumination (Figure 2A). *f* decreased by 1.2 and 2.0 Hz, respectively (t-test, p = 0.001 and p = 0.0001, n = 6). Apnea – no inspiratory effort – resulted in more than one-third of all trials at 10.2 mW (i.e., 11 of 30 bouts, e.g., Figure 2A, bottom). V_T_ decreased in response to 8.6 and 10.2 mW light in both cases by 0.03 ml (t-test, p = 0.04 and p = 0.02, n = 6). MV decreased by 11 and 20 ml/min, respectively (t-test, both p = 0.02, n = 6) (Figure 2B).

In comparison, sedated wild-type littermates subjected to the same protocol showed no light-evoked changes in breathing (Figure 2 – figure supplement 1A and 1B).

We repeated these experiments in Dbx1;ArchT mice while awake and unrestrained (Figure 2C). The lowest intensity light (6.8 mW) decreased *f* and V_T_ by 0.01 Hz and 0.03 ml, respectively (neither change was statistically significant by t-test, p = 0.06 and 0.07, n = 5). MV decreased significantly by 7.4 ml/min (t-test, p = 0.04, n = 5) (Figure 2D).

The effects on breathing were more profound when we illuminated at 8.6 and 10.2 mW (Figure 2C). *f* decreased by 1.1 and 1.2 Hz, respectively (t-test, p = 0.002 and p = 0.02, n = 5) and MV decreased by 22 and 32 ml/min, respectively (t-test, p = 0.04 and p = 0.03, n = 5). One animal stopped breathing for ~4 s (i.e., apnea, Figure 2C, bottom trace). Although V_T_ decreased by 0.05 and 0.15 ml, respectively, statistical hypothesis testing did not detect significant light-induced changes (t-test, p = 0.2 and p = 0.08, n = 5), probably due to the high variability of V_T_ in awake animals (Figure 2D).

In comparison, awake unrestrained wild-type littermates showed no changes in breathing in reponse to light of any intensity (Figure 2 – figure supplement 1C and 1D).

Therefore, these data collectively show that ArchT-mediated Dbx1 preBötC neuron hyperpolarization reduces breathing up to and including apnea in sedated and awake intact mice.

Next we applied brief (100 ms) light pulses randomly during the breathing cycle, which we defined as spanning 0-360° (see Materials and Methods, Figure 3 inset). Brief photoinhibition of the preBötC early during inspiration (Φ_Stim_ of 0-30°) caused a phase advance such that the subsequent inspiration occurred earlier than expected (Φ_Shift_ = -147 ± 23°, p = 1e-6, n = 4) while shortening inspiratory time (T_i_) by almost half (ΔT_i_ = 45 ± 5%, p = 1e-6, n = 4) (Figure 3A_1,2_ and A_3_ top trace). Brief photoinhibition also evoked significant phase advances and reduced T_i_ during the rest of inspiration (Φ_Stim_ of 30-120°), but the magnitude of those changes monotonically decreased as Φ_Stim_ approached the inspiratory-expiratory transition.

**Figure 3.**
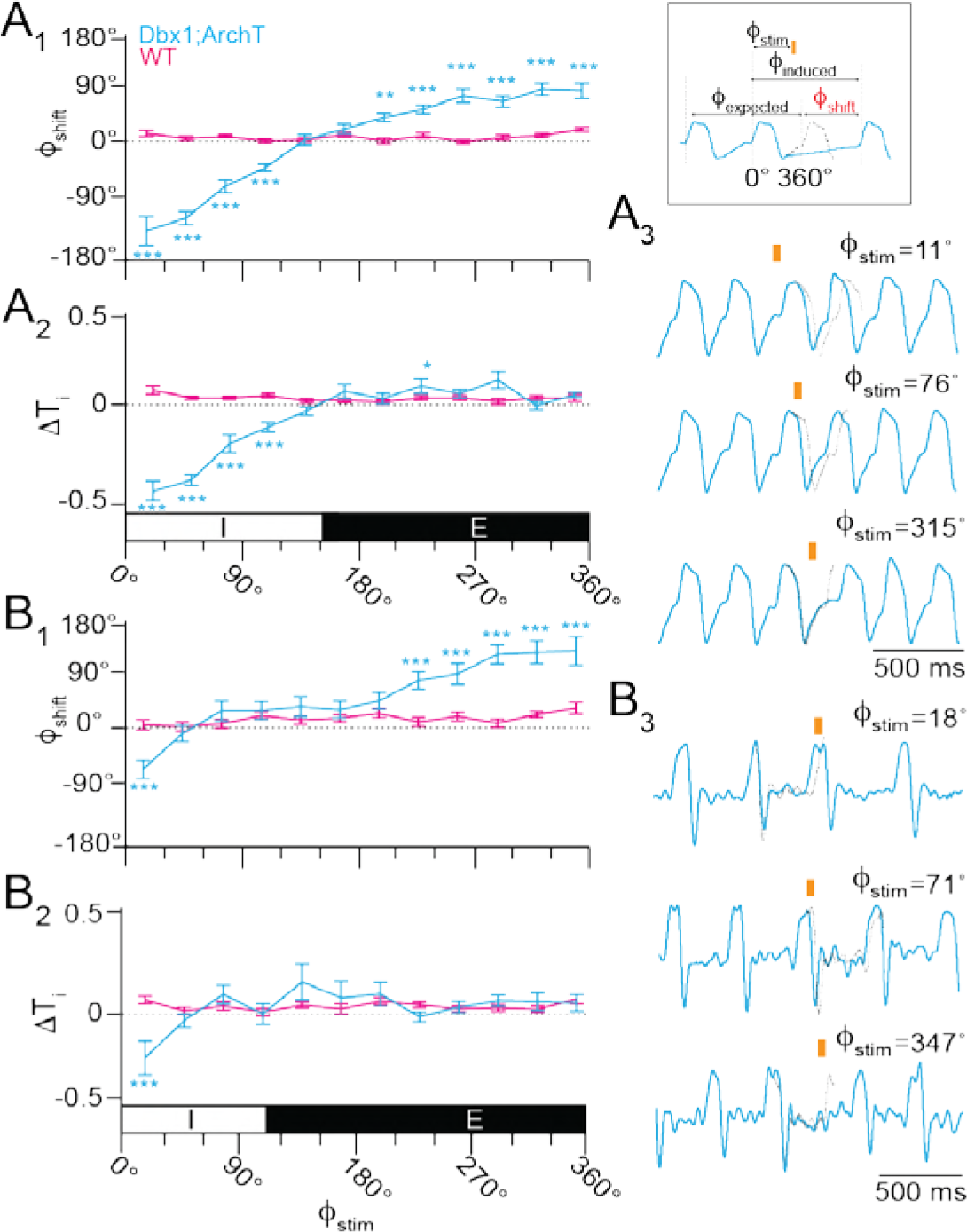
Effects of brief photoinhibition on the breathing phase and inspiratory duration in Dbx1;ArchT mice (n = 6 in A, n = 5 in B, cyan) and wild type littermates (n = 6, magenta). **A_1_**, Phase-response curve plotting Φ_Shift_ following 100-ms photoinhibition at Φ_Stim_ throughout the breathing cycle in sedated mice. Φ_Stim_ was partitioned into 12 equally sized bins (30°) in A and B. **A_2_**, Phase-response curve showing changes in T_i_ following brief photoinhibition (i.e., the perturbed breath) in the same cohort of sedated mice. The abscissa marks the inspiratory (I, 0-150°) and expiratory (E, 150-360°) phases of the breathing cycle (0-360°), which applies to A_1_ and A_2_. **A_3_**, Sample airflow traces from a representative sedated mouse (Φ_Stim_ is indicated by an orange bar and numeral value). Time calibration is shown. **B_1_**, Phase-response curve plotting Φ_Shift_ following brief photoinhibition at Φ_Stim_ throughout the breathing cycle in awake unrestrained mice. **B_2_**, Phase-response curve showing changes in T_i_ following brief photoinhibition (i.e., the perturbed breath) in the same cohort of awake unrestrained mice. The abscissa marks the inspiratory (I, 0-110°) and expiratory (E, 110-360°) phases of the breathing cycle (0-360°), which applies to B_1_ and B_2_. **B_3_**, Sample airflow traces from a representative awake unrestrained mouse (Φ_Stim_ is indicated by an orange bar and numeral value). Time calibration is shown.

Brief photoinhibition did not perturb the system during the inspiratory-expiratory transition (Φ_Stim_ of 120-180°). During early expiration (Φ_Stim_ of 180-210°), which is often referred to as post-inspiration (Anderson et al., 2016; Dutschmann et al., 2014) we observed the first significant phase delay such that the subsequent inspiration occurred later than expected in response to brief photoinhibition (Φ_Shift_ = 32 ± 7°, p = 0.006, n = 4, Figure 3A_1_ and A_3_ bottom trace). Phase delays were consistently evoked during expiration (Φ_Stim_ of 210-360°) with a maximum phase delay during late expiration (Φ_Stim_ of 300-330°) (Φ_Shift_ = 78 ± 10°, p = 1e-6, n = 4). Brief photoinhibition during expiration did not affect T_i_, which is a straightforward result because the inspiratory period had ended (Figure 3A_2_). Note, that ΔT_i_ was statistically significant at Φ_Stim_ of 210-240°) but that change is not physiologically meaningful because the magnitude of the change is small and not part of a consistent trend in the phase-response curve.

The relationship between Φ_Stim_ and the phase of the subsequent breath (Φ_N+1_, Figure 3 – figure supplement 1A_1_) closely resembled the relationship between Φ_Stim_ and Φ_Shift_ (Figure 3A_1_), which suggests that brief photoinhibition resets the phase of the oscillator.

In contrast to its effects on breathing phase (Φ_Shift_ and Φ_N+1_), brief photoinhibition had little effect on V_T_ throughout most of the respiratory cycle with changes of less than 10% across the entire respiratory cycle, except during early inspiration (Φ_Stim_ of 0-30°, in which V_T_ decreased by 23 ± 8%, p = 0.02, n = 4) and early expiration (Φ_Stim_ of 150-180°, in which V_T_ increased by 16 ± 11%, p = 0.01, n = 4) (Figure 3 – figure supplement 1A_2_). Despite the fact that two out of 12 measurements pass the threshold for statistical significance, these data do not convincingly demonstrate that brief photoinhibition of Dbx1 preBötC neurons systematically influences V_T_ in sedated mice.

We repeated brief photoinhibition experiments in awake unrestrained Dbx1;ArchT mice. The plots of Φ_Shift_, ΔT_i_, Φ_N+1_, and ΔV_T_ versus Φ_Stim_ were qualitatively similar to the experiments in sedated mice (compare Figure 3A to 3B and Figure 3 – figure supplement 1A to 1B).

Photoinhibition during early inspiration (Φ_Stim_ of 0-30°) caused a phase advance (Φ_Shift_ = -86 ± 16°, p = 1e-5, n = 4). The first significant phase delay in the awake animal occurred when brief photoinhibition was applied during peak expiration (Φ_Stim_ of 210-240°, Φ_Shift_ = 68 ± 15°, p = 1e-6, n = 4). Φ_Shift_ tended to increase as brief photoinhibition was applied at later points during the expiratory phase. The maximum phase delay occurred during late expiration (Φ_Stim_ of 330-360°, Φ_Shift_ = 118 ± 25°, p = 4e-5, n = 4) (Figure 3B_1_ and B_3_). Brief photoinhibition decreased T_i_ by nearly one-third (ΔT_i_ = 28 ± 9%, p = 1e-5, n = 4) during early inspiration (Φ_Stim_ of 0-30°) but had no significant effect at any other time during the cycle.

### Photostimulation of Dbx1 preBötC neurons enhances breathing and modifies the timing and magnitude of breaths

We illuminated the preBötC unilaterally in sedated adult Dbx1;CatCh mice (the intersection of a *Dbx1*^CreERT2^ driver mouse and a reporter featuring Cre- and Flp-dependent calcium translocating channelrhodopsin [CatCh] expression) following viral transduction in the preBötC with a synapsin-driven Flp recombinase. Using this double-stop intersectional approach, CatCh-EYFP expression was limited to the preBötC (Figure 1D). In control conditions *f* was typically ~3 Hz, V_T_ was ~0.1 ml, and MV was ~50 ml/min. Bouts of blue light (473 nm) at three intensities significantly increased *f* by 0.8, 1.1, and 1.3 Hz, respectively (t-test, p = 0.03, 0.005, and 0.03, n = 4). There were no significant effects on V_T_ or MV at any light intensity (Figure 4A and B).

**Figure 4.**
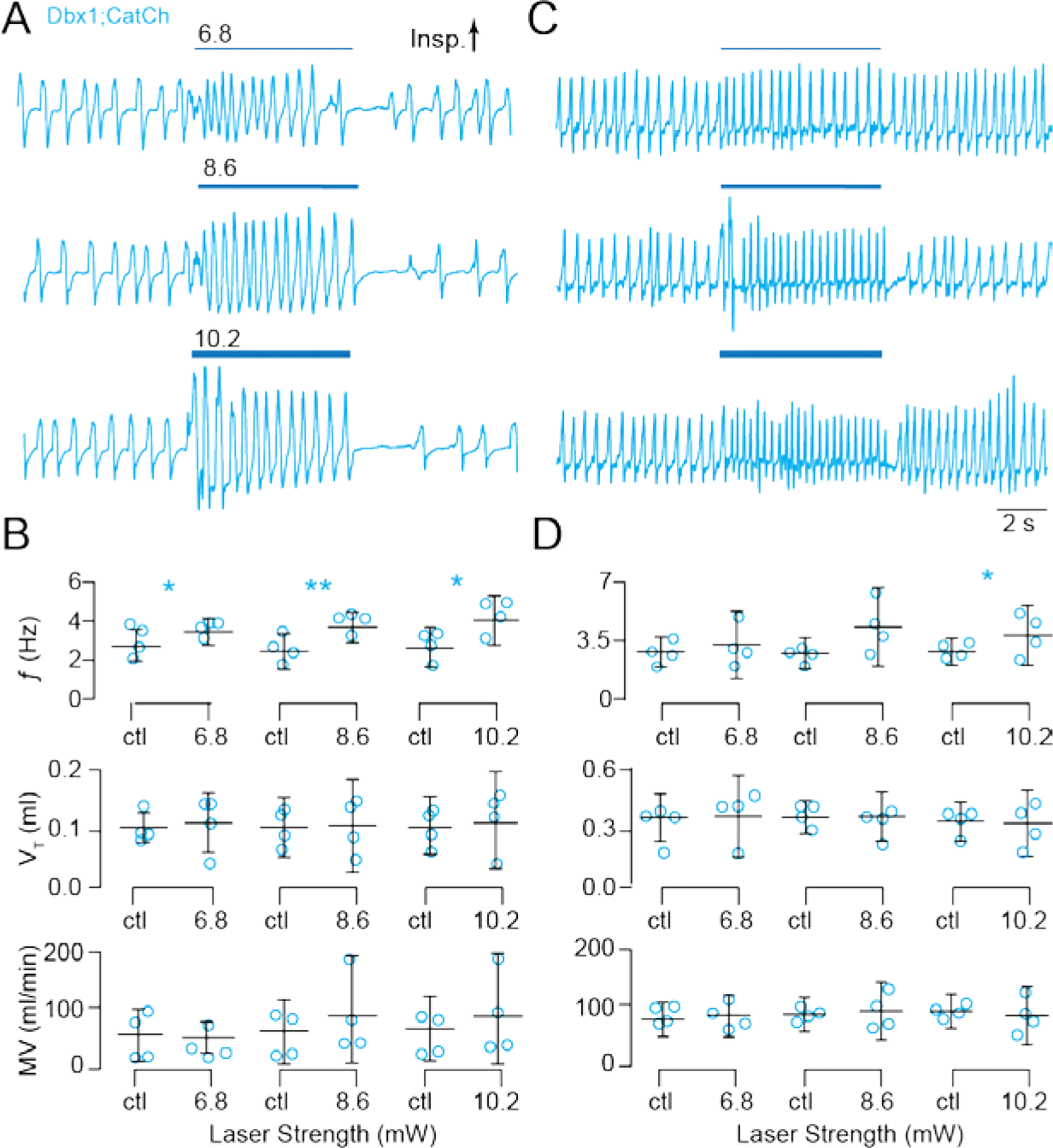
Photostimulation of Dbx1 preBötC neurons speeds-up breathing in adult Dbx1;CatCh mice. **A**, Airflow traces from a sedated mouse exposed to 5-s bouts of unilateral preBötC illumination at three intensities (units of mW). Cyan line thickness corresponds to light intensity, which is also annotated above each line. **B**, Group data from experiments in A quantifying light-evoked changes in *f*, V_T_ and MV. Symbols show the mean *f*, V_T_, and MV measured in each mouse. Bars show the mean and SD for all animals tested (n = 4). Control measurements are labeled ‘ctl’; numerals indicate light intensity. **C**, Airflow traces from an awake unrestrained mouse exposed to 5-s bouts of bilateral preBötC illumination at three intensities. Cyan line thickness corresponds to light intensity; annotations mach those in A. **D**, Group data from experiments in C quantifying light-evoked changes in *f*, V_T_ and MV. Symbols show the mean *f*, V_T_, and MV measured in each mouse. Bars show the mean and SD for all animals tested (n = 4). Control measurements are labeled ‘ctl’; numerals indicate light intensity. Asterisks represent statistical significance at p < 0.05; the double asterisk represents p < 0.01.

We repeated these unilateral photostimulation experiments in Dbx1;CatCh mice while awake and unrestrained. Frequency increased by 1.6 Hz in response to light at the highest intensity (Figure 4C and D). There were no other notable changes in *f*, V_T_, or MV at any light intensity.

In wild type littermates, we observed no effects on breathing in either sedated or awake mice in response to light at any intensity (Figure 4 – figure supplement 1).

Therefore, these data collectively show that CatCh-mediated photostimulation of Dbx1 preBötC neurons selectively enhances breathing frequency in sedated and awake intact mice.

Next we applied brief (100 ms) light pulses at different time points during the breathing cycle. Unilateral illumination of the preBötC during inspiration caused a phase delay and increased T_i_. The maximum phase delay occurred during peak inspiration (Φ_Stim_ of 60-90°, Φ_Shift_ = 125 ± 18°, p = 1e-6, n = 4) (Figure 5A_1_) and coincided with the maximum ΔT_i_ (29 ± 7%, p = 1e-6, n = 4) (Figure 5A_2_). Brief photostimulation caused a phase advance during the inspiratory-expiratory transition (Φ_Stim_ of 90-120°) and throughout expiration (Φ_Stim_ ≥ 120°) without affecting T_i_. The maximum phase advance occurred during early expiration (Φ_Stim_ of 150-180°, Φ_Shift_ = -128 ± 4°, p = 1e-6, n = 4) (Figure 5A_1_ and A_3_). The relationship between Φ_Stim_ and the phase of the subsequent breath (Φ_N+1_, Figure 5 – figure supplement 1A_1_) mimicked the relationship between Φ_Stim_ and Φ_Shift_ (Figure 5A_1_), which suggests that brief photostimulation resets the phase of the oscillator. We observed no effects of brief photostimulation on V_T_ (Figure 5 – figure supplement 1A_2_).

**Figure 5.**
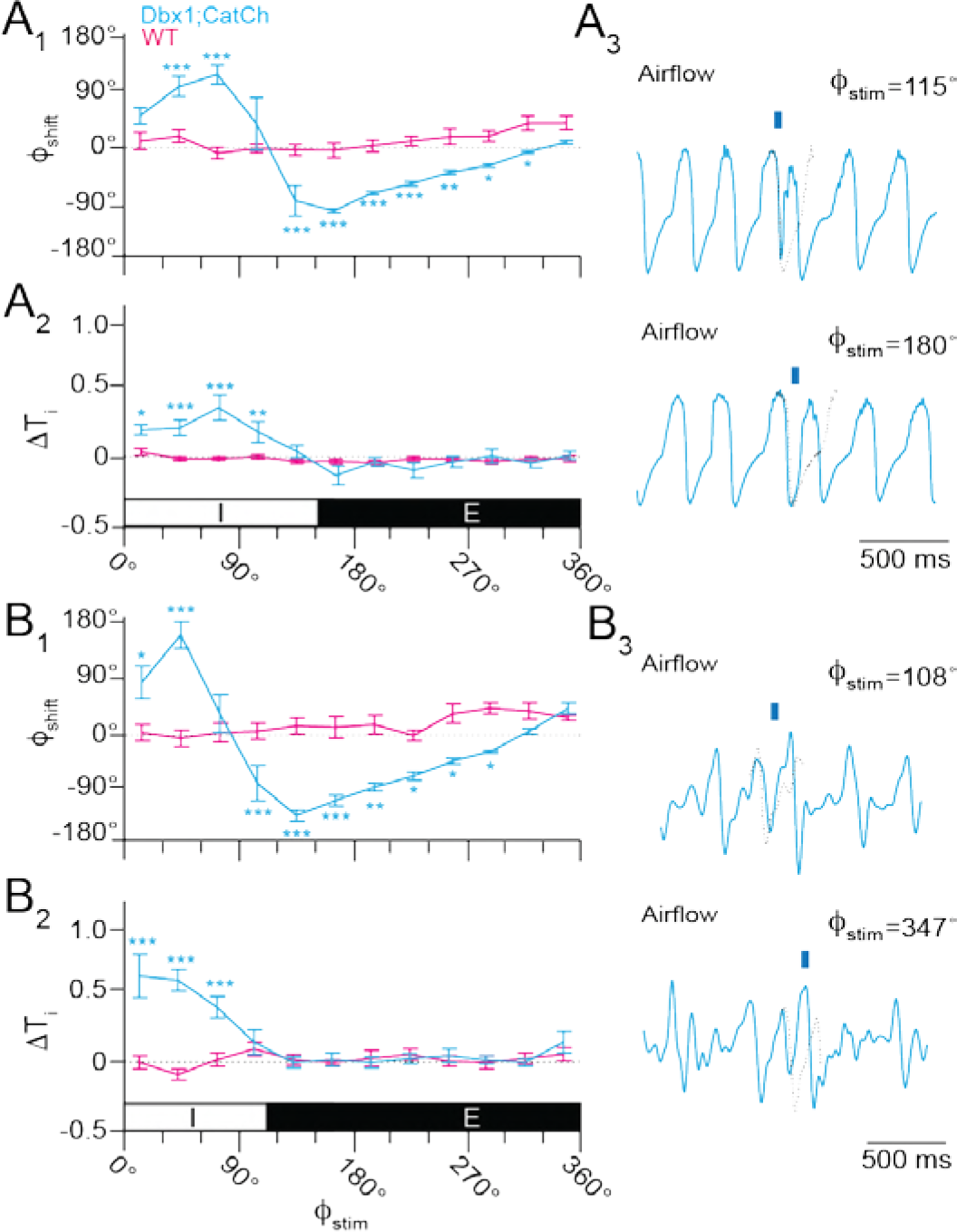
Effects of brief photostimulation on the breathing phase and inspiratory duration from Dbx1;CatCh mice (n = 4, cyan) and wild type littermates (n = 4, magenta). **A_1_**, Phase-response curve plotting Φ_Shift_ following 100-ms photostimulation at Φ_Stim_ throughout the breathing cycle in sedated mice. Φ_Stim_ was partitioned into 12 equally sized bins (30°) in A and B. **A_2_**, Phase-response curve for changes in T_i_ following photostimulation (i.e., the perturbed breath) in the same cohort of sedated mice. The abscissa marks the inspiratory (I, 0-150°) and expiratory (E, 150-360°) phases of the breathing cycle (0-360°), which applies to A_1_ and A_2_. **A_3_**, Sample airflow traces from a representative sedated mouse (Φ_Stim_ is indicated by an orange bar and numeral value). Time calibration as shown. **B_1_**, Phase-response curve plotting Φ_Shift_ following brief photostimulation at Φ_Stim_ throughout the breathing cycle in awake unrestrained mice. **B_2_**, Phase-response curve for changes in T_i_ following brief photostimulation (i.e., the perturbed breath) in the same cohort of awake unrestrained mice. The abscissa marks the inspiratory (I, 0-110°) and expiratory (E, 110-360°) phases of the complete breathing cycle (0-360°), which applies to B_1_ and B_2_. **B_3_**, Sample airflow traces from a representative awake unrestrained mouse (Φ_Stim_ is indicated by an orange bar and numeral value). Time calibration is shown.

We repeated brief photostimulation experiments in awake intact Dbx1;CatCh mice. The plots of Φ_Shift_ and ΔT_i_ vs Φ_Stim_ were qualitatively similar to those recorded in sedated mice (compare Figure 5B to 5A). Brief photostimulation during early and mid-inspiration (Φ_Stim_ of 0-60°) caused a phase delay (maximum Φ_Shift_ = 147 ± 52, p = 1e-5, n = 4) (Figure 5B_1_). We measured no phase shift for late inspiration (Φ_Stim_ of 60-90°). The phasic effect of brief photostimulation changed sign around the inspiratory-expiratory transition (Φ_Stim_ ≥ 90°); brief photostimulation subsequently evoked breaths earlier than expected. We measured the maximum phase advance during early expiration (Φ_Stim_ of 120-150°, Φ_Shift_ = -159 ± 9°, p = 1e-5, n = 4) (Figure 5B_1_). The last statistically significant phase delay occurred during late expiration (Φ_Stim_ of 270-300°, Φ_Shift_ = -52 ± 3°, p = 0.05, n = 4).

Brief photostimulation of Dbx1 preBötC neurons in awake intact mice also extended T_i_ during inspiration (Figure 5B_2_); the effect was even more pronounced than in sedated mice (Figure 5A_2_). The maximum ΔT_i_ occurred during early inspiration (Φ_Stim_ of 0-30°) in which T_i_ increased by over half (56 ± 14%, p = 1e-6, n = 4). The ability of brief photostimulation to extend T_i_ decreased during the inspiratory phase (Figure 5B_1_) such that no significant effects occurred after Φ_Stim_ exceeded 90°. The relationship between Φ_Stim_ and Φ_N+1_ illustrated a phase delay evoked by brief photostimulation during mid-inspiration (Φ_Stim_ of 30-60°, Figure 5 – figure supplement 1A_1_), which partially recaps the relationship that was more pronounced in the plot of Φ_Shift_ vs. Φ_Stim_ (Figure 5A_1_). We observed no relationship for ΔV_T_ vs. Φ_Stim_ (Figure 5 – figure supplement 1B_2_), as in the sedated mouse (Figure 5 – figure supplemnt 1B_1_).

These data are consistent photostimulus-induced resetting of the inspiratory oscillator, although the data are noisier in the awake adult, freely behaving mouse.

## DISCUSSION

### Role diversity challenges the Dbx1 core hypothesis

The idea that Dbx1 preBötC neurons are inspiratory rhythmogenic has become generally well accepted, but it must be reevaluated given the expanding spectrum of non-rhythmogenic and non-respiratory functions attributed to this neuron class, particularly in adult animals.

Perinatally Dbx1 preBötC neurons generate rhythm and pattern. *Dbx1* knock-out mice do not breathe and form no recognizable preBötC (Bouvier et al., 2010; Gray et al., 2010), the site of inspiratory rhythmogenesis (Del Negro et al., 2018; Feldman and Del Negro, 2006; Feldman et al., 2013; Ramirez et al., 2016; Smith et al., 1991). Their selective destruction in a slice model of breathing (Funk and Greer, 2013) slows and then stops the rhythm, evidence of their rhythmogenic role, while also attenuating airway-related XII motor output (Wang et al., 2014) because of Dbx1 premotor neurons in the preBötC that drive XII (Revill et al., 2015; Wang et al., 2014) as well as phrenic motoneurons (Wu et al., 2017).

This theme continues in adult mice. Sst-expressing preBötC neurons, ~17% of the Dbx1-derived population, appear to lack rhythmogenic function but rather shape motor output pattern (Cui et al.,2016), q.v., (Koizumi et al., 2016). More than half (56%) of Dbx1 preBötC neurons characterized by Cdh9 expression lack respiratory rhythmicity but project to the locus coeruleus and putatively influence arousal (Yackle et al., 2017). If we assume that non-Sst and non-Cdh9 Dbx1 neurons have respiratory functions, and that individual neurons do not fulfill multiple duties, then these statistics suggest that not more than 27% of Dbx1 preBötC neurons in adult mice are exclusively rhythmogenic.

### Photoinhibition and photostimulation demonstrate Dbx1 preBötC neurons influence rhythm and pattern

Sustained photoinhibition caused graded frequency decreases including apnea, which are evidence that Dbx1 neurons form the core oscillator. However, photoinhibition also decreased V_T_, indicating that Dbx1 neurons also govern breath size, i.e., motor pattern. We reported qualitatively similar data in (Vann et al., 2016) but the effects were more mild because of the weaker archaerhodopsin variant available at the time. Dbx1 neurons that influence airway and pump-related motor function have been analyzed in detail (Cui et al.,2016; Revill et al., 2015; Wang et al., 2014; Wu et al., 2017). Here we limit our comments to acknowledging those motor-related roles, and we concentrate on analyzing the role of Dbx1 preBötC neurons in rhythmogenesis.

Sustained photostimulation approximately doubled the breathing rate from ~3.5 to 7 Hz. In contrast, Baertsch and colleagues (Baertsch et al., 2018) reported minor (~10%) frequency changes in vagus intact mice in response to sustained photostimulation. These two results are not discrepant, even if they appear to be at face value. We were able to evoke higher frequencies in our experiments most likely due to the accelerated response time, enhanced light sensitivity, larger voltage responses evoked by photoactivated CatCh compared to ChR2 (Kleinlogel et al., 2011), and the fact that we applied laser strengths up to 10.2 mW whereas Baertsch *et al.* purposely limited their pulses to 7 mW or less (Baertsch et al., 2018). Those authors showed that phasic synaptic inhibition critically influences breathing frequency and we do not disagree. We purposely did not vagotomize our mice to preserve phasic synaptic inhibition and thus high breathing frequencies are possible during photostimulation.

### Phase-response experiments demonstrate that Dbx1 preBötC neurons are rhythmogenic

If Dbx1 preBötC neurons are inspiratory rhythmogenic, then transiently stimulating them should evoke inspiratory breaths at any point in the breathing cycle except, potentially, during the post-inspiratory (early expiratory) refractory period identified *in vitro* (Guerrier et al., 2015; Kottick and Del Negro, 2015) and in vagotomized mice *in vivo* (Baertsch et al., 2018). We evoked inspiratory breaths at all points during the respiratory cycle without evidence of a refractory period. Brief photostimulation during inspiration prolonged it (i.e., increased T_i_) and delayed the next cycle (i.e., a phase delay). The straightforward interpretation is that CatCh-mediated inward current augments recurrent excitation thus prolonging inspiratory burst duration. Overexcited rhythmogenic neurons require more time to recover, which lengthens cycle time and delays the subsequent inspiration.

We observed that photostimulation at any other point in the cycle evoked inspiration earlier than expected, a phase advance, but did not otherwise modify inspiration. In contrast to a prior report, brief photostimulation did not evoke phase advances during early expiration (Alsahafi et al., 2015). But in that experimental context a synapsin promoter drove channelrhodopsin expression in both excitatory and inhibitory preBötC neurons. Because preBötC rhythmogenesis depends on recurrent excitation, and the network is at the nadir of its excitability during early expiration (Del Negro et al., 2018; Feldman and Kam, 2015; Ramirez et al., 2016), photostimulation of inhibitory neurons in concert with excitatory neurons would be less effective to evoke inspiratory bursts during early expiration.

Selective photostimulation of excitatory Dbx1-derived preBötC neurons should evoke phase advances during early expiration, and it does. Cui *et al*. (2016) photostimulated excitatory Dbx1 neurons and evoked phase advances of up to ~72° during most of expiratory phase, except during the inspiratory-expiratory transition. We evoked more substantial phase advances of 90-150° during the early expiration. These results are not in conflict, but key methodological differences may explain the discrepancy. Cui *et al.* anesthetized their mice and applied a maximum laser power of 7 mW to activate channelrhodopsin, whereas we used awake or lightly sedated mice and applied a maximum laser power of 10.2 mW to activate the channelrhodopsin variant CatCh. Assuming that the fiberoptic appliances in both studies equally attenuate laser power from box to preBötC, then the larger phase advances we evoked during early expiration could be attributable to a higher excitability level of the preBötC in the unanesthetized (or lightly sedated) mice, higher laser power, as well as the accelerated response time, enhanced light sensitivity, and larger voltage responses evoked by photoactivated CatCh compared to ChR2 (Kleinlogel et al., 2011).

Brief photoinhibition of Dbx1 preBötC neurons during inspiration shortened it (i.e., decreased T_i_) and initiated the next cycle earlier than expected, a phase advance. We infer that hyperpolarizing rhythmogenic neurons checks the recurrent excitation process, which impedes but does not prevent inspiration. Nevertheless, the evoked breath is shorter in duration. preBötC neurons do not overexcite or become refractory, which facilitates the onset of the next cycle, hence the phase advance. That mechanism, here evoked by ArchT, mirrors the role of endogenous phasic synaptic inhibition, which curbs recurrent excitation to limiting inspiratory activity and facilitate inspiratory-expiratory phase transition (Baertsch et al., 2018). We found that photoinhibition during expiration consistently caused a phase delay, which indicates hyperpolarization of Dbx1 preBötC neurons resets recurrent excitation and thus prolongs the interval until the next inspiration.

Our interpretations of the phase-response experiments, both photostimulation and photoinhibition, are consistent with Dbx1 preBötC neurons having direct temporal control over inspiration as well as post-inspiration and the expiratory interval. That conclusion may seem overly broad considering, first, that the preBötC is the acknowledged inspiratory oscillator and, second, that oscillator microcircuits for post-inspiration (the postinspiratory complex, PiCo, Anderson et al., 2016) and expiration (the lateral parafacial group, pF_L_, Huckstepp et al., 2016, 2015; Pagliardini et al., 2011) also exist. Nevertheless, the preBötC plays a dominant role in organizing all phases of breathing by entraining the other oscillators in intact mice, and in reduced preparations that retain PiCo and pF_L_ (Del Negro et al., 2018; Moore et al., 2013; Ramirez et al., 2016). Therefore, the present data are consistent with Dbx1 preBötC interneurons constituting the oscillator core for inspiration and the central organizer for breathing.

### Could optogenetic perturbation of inputs to the preBötC modulate breathing?

The intersectional mouse genetics in Dbx1;ArchT mice leads to fusion protein expression in Dbx1-derived cells throughout the neuraxis. Therefore, preBötC illumination inhibits constituent interneurons but also axons of passage and the axon terminals of Dbx1 neurons from remote locations (Ruangkittisakul et al., 2014) that could disfacilitate the preBötC. If disfacilitation were primarily modulating preBötC activity in Dbx1;ArchT mice, then light-evoked hyperpolarization should be commensurate in non-Dbx1 neurons (which do not express ArchT) and Dbx1 neurons; and, TTX should block it in both cases. However, non-Dbx1 neurons hyperpolarized ~1 mV in response to maximum illumination whereas Dbx1 neurons hyperpolarized ~11 mV, and TTX did not notably affect either response. We conclude that direct postsynaptic hyperpolarization of Dbx1 preBötC neurons, rather than a reduction of tonic excitatory drive, is the predominant effect of preBötC illumination in Dbx1;ArchT mice.

Light-evoked breathing changes in Dbx1;CatCh mice cannot be explained by photostimulation of axon terminals and axons of passage that originate outside of, but synapse within, the preBötC. We used double-stop technology to limit CatCh expression to Dbx1-derived neurons (not glia, see below), whose somas reside in the preBötC or directly adjacent sites including the Bötzinger complex of inhibitory neurons (Ezure et al., 2003; Tanaka et al., 2003), and the rostral ventral respiratory group (Dobbins and Feldman, 1994; Ellenberger and Feldman, 1990; Gaytán et al., 2002) of excitatory phrenic premotor neurons. If Dbx1-derived expiratory neurons in the Bötzinger complex exist (which has not been demonstrated), then their photostimulation would depress breathing (Janczewski et al., 2013; Marchenko et al., 2016), the opposite of what we measured. If photostimulation affected Dbx1 phrenic premotor neurons in the rostral ventral respiratory group (Wu et al., 2017), then that would enhance the magnitude of inspiratory breaths, but not the inspiratory timing circuits in the preBötC. Sustained photostimulation experiments only enhanced breathing frequency and never V_T_, which diminishes the likelihood that our protocols influenced Dbx1-derived phrenic premotoneurons. Thus, this caveat is unlikely to affect our primary conclusions regarding rhythmogenesis.

### Effects on Dbx1-derived glia in the preBötC

Dbx1-expressing precursor cells develop into neurons and glia (Bouvier et al., 2010; Gray et al., 2010; Kottick et al., 2017; Ruangkittisakul et al., 2014) but optogenetic perturbation of glia is unlikely to have influenced the present results. First, we consider photoinhibition. Astrocytes support excitatory synaptic function in the preBötC (Hülsmann et al., 2000), but that role is metabolic in nature and light-evoked hyperpolarization would not preclude it. Calcium excitability and gliotransmission, which could be affected by photoinhibition, pertain to purinergic modulation and hypoxic challenges to the preBötC (Angelova et al., 2015; Funk et al., 2015; Huxtable et al., 2010; Rajani et al., 2017), but are less relevant factors governing the basal breathing state, which is the baseline for our experiments.

Photostimulation experiments unambiguously identify neurons as the cellular population that forms the core inspiratory oscillator. CatCh expression was induced following Cre/Lox and Frt/Flp recombination. We used a synapsin promoter to express Flp locally in the preBötC so only Dbx1 neurons would be transfected and express CatCh.

ArchT expression is selectively (but not exclusively) limited to neurons by the timing of tamoxifen administration. Inducing Cre/lox recombination in pregnant *Dbx1*^CreERT2^ mice at E9.5 reduces ArchT expression in glia to ~40%, whereas ArchT expression in neurons remains above 90% (Kottick et al., 2017), which increases our confidence that photoinhibition largely affects neurons (not glia) and that neurons are the predominate rhythmogenic constituents and most parsimonious explanation for the light-induced changes in breathing.

### Size of the Dbx1 core oscillator

Up to 73% of Dbx1 preBötC neurons serve non-rhythmogenic functions: 56% influence arousal (Yackle et al., 2017) and 17% influence motor pattern (Cui et al., 2016), which accounts nearly three-quarters of the Dbx1 population in the preBötC. What implications does that have for the composition and size of the inspiratory core oscillator whose constituent interneurons are Dbx1-derived too?

Dbx1-Cdh9 preBötC neurons were certainly photoinhibited and photostimulated in our experiments. However, those neurons influence behavioral state (e.g., eupnea, grooming, exploring, sniffing, etc.) rather than cycle-to-cycle breathing dynamics. We applied optogenetic perturbations only during eupnea, not during grooming or active movement, to control for behavioral shifts. Given that Dbx1-Cdh9 neurons are either weakly or not rhythmic (Yackle et al., 2017), briefly perturbing them would not influence the phase-response relationships, and thus would not confound our interpretation that Dbx1 preBötC neurons (even if a limited fraction of them) comprise the core oscillator.

Illumination of Sst-expressing Dbx1 neurons could be responsible for the decreases in V_T_ and apneas we report during sustained photoinhibition. In general, perturbations of Sst-expressing preBötC neurons affect breathing motor pattern in vagotomized and non-vagotomized adult mice (Cui et al.,2016; Koizumi et al., 2016); those effects are strong enough to completely stop breathing movements in intact adult rats (Tan et al., 2008). Our experiments would only impact neurons that are both Dbx1-dervied and Sst-expressing, thus a smaller population than Tan *et al.* (2008) manipulated. Nevertheless, to the extent that photoinhibition decreased breath magntidue and caused apnea, we attribute in part to direct effects on pattern-related Sst-expressing Dbx1-derived preBötC neurons that are either premotor part of a larger pattern-generating system (Cui et al.,2016; Revill et al., 2015; Wu et al., 2017).

If Cdh9 and Sst subpopulations of Dbx1 preBötC neurons are independent of the core respiratory oscillator, then only a small fraction (~27%) of Dbx1 neurons are available for rhythmogenesis. Dbx1 neurons that comprise the preBötC core number approximately 600 (Kottick et al., 2017; Wang et al., 2014). If one excludes Cdh9 and Sst neurons from this estimation, then as few as 160 Dbx1 preBötC neurons would remain for rhythmogenesis (we assue subpopulations serve one function). Can such a small number of interneurons comprise the inspiratory core oscillator?

Holographic photolysis of caged glutamate onto 4-9 preBötC neurons evokes inspiratory motor output *in vitro* (Kam et al., 2013). This type of stimulation would affect Dbx1-Cdh9 neurons that are weakly or non-rhythmic (Kam et al., 2013; Yackle et al., 2017) as well as inhibitory preBötC neurons (Kuwana et al., 2006; Morgado-Valle et al., 2010; Winter et al., 2009) so it may overestimate the minimum number of activated preBötC neurons needed to evoke inspiratory bursts. Regardless, a reasonable conclusion is that stimulating relatively small numbers of preBötC neurons are capable of inducing inspiratory burst cycles, which lends credence to the notion that a small subfraction of Dbx1 preBötC neurons could be rhythmogenic in the midst of a potentially larger population of non-rhythmogenic (both pattern-generating and non-respiratory) preBötC neurons.

Glutamatergic preBötC neurons not derived from Dbx1-expressing precursors may also comprise part of the core oscillator (Baertsch et al., 2018; Koizumi et al., 2016). We cannot precisely estimate the size of that subpopulation but we expect that it will be small based on the small fraction of preBötC neurons that express Vglut2 but not Dbx1 (Bouvier et al., 2010; Gray et al., 2010).

### Dbx1 core hypothesis

The rhythmogenic subset of Dbx1 preBötC interneurons may be small, perhaps as little as 27% of the total Dbx1 population, but their outsize contribution to rhythmogenesis is unmistakable given the robust effects of sustained and transient photoinhibition and photostimulation demonstrated here, and by prior reports (Alsahafi et al., 2015; Cui et al.,2016; Koizumi et al., 2016). Therefore, whatever else Dbx1 preBötC neurons do – influence motor pattern and behavioral state – they certainly comprise the inspiratory core oscillator. Two key challenges going forward will be, first, to quantify the proportion of the rhythmogenic preBötC core that is non-Dbx1-derived, and second, to discriminate either on the basis of genetic or other markers, rhythmogenic from non-rhythmogenic Dbx1 neurons.

## MATERIALS AND METHODS

### Mice

The Institutional Animal Care and Use Committee at The College of William and Mary approved these protocols. Female mice that express tamoxifen-sensitive Cre recombinase in *Dbx1*-derived progenitor cells, i.e., *Dbx1*^CreERT2^ (Ruangkittisakul et al., 2014), available at Jax (strain 028131, Jackson Laboratories, Bar Harbor, ME, USA), were mated with males from two different reporter strains. The first reporter strain expresses an Archaerhodopsin-3 tagged with EGFP fusion protein (ArchT-EGFP) in a Cre-dependent manner from the endogenous *Gt(ROSA)26Sor* locus (Allen Institute nomenclature, Ai40D; Jax strain #021188,). The second reporter strain features *Frt*- and *LoxP*-flanked STOP cassettes followed by a fusion gene coding for calcium translocating channelrhodopsin and EYFP (CatCh-EYFP), which is expressed following Cre- and Flp-mediated recombination (Allen Institute nomenclature, Ai80D; Jax strain #025109). We administered tamoxifen to pregnant dams (22.5 mg/kg) at embryonic day 9.5 to maximize neuronal expression and minimize glial expression (Kottick et al., 2017). Dbx1;ArchT or Dbx1;CatCh mice were distinguished from wildtype (WT) littermates, which lack EGFP or EYFP, via post-hoc histology. Therefore, WT littermates formed a control group whose constituent members were unknown to the experimenter.

### Brainstem slices

Neonatal Dbx1;ArchT mice (0-4 days old) were anesthetized via hypothermia, decerebrated, and then dissected in 4° C artificial cerebrospinal fluid (aCSF) containing (in mM): 124 NaCl, 3 KCl, 1.5 CaCl_2_, 1 MgSO_4_, 25 NaHCO_3_, 0.5 NaH_2_PO_4_, and 30 dextrose aerated continually with carbogen (95% O_2_ and 5% CO_2_) at pH 7.4. The isolated neuraxes were glued to an agar block and mounted rostral side up in the vise of a vibratome. We cut the neuraxes in the transverse plane to obtain a single 500-μm-thick section containing the preBötC as well as the hypoglossal (XII) cranial motor nucleus and its rostral nerve rootlets. The anatomical criteria for isolating the preBötC in rhythmically active slices from neonatal Dbx1-reporter mice are detailed in a series of open access atlases (Ruangkittisakul et al., 2014). Slices were anchored using a silver wire grid in a recording chamber on a fixed-stage upright physiology microscope. We perfused the slice with aCSF at 27° C (2 ml/min) and elevated the K^+^ concentration to 9 mM. Inspiratory motor output was recorded from the XII nerve rootlets using a differential amplifier (gain 2000x) and a band-pass filter (300-1000 Hz). Nerve root output was full-wave rectified and smoothed for display.

We identified Dbx1 neurons under epifluorescence via EGFP expression and then performed whole-cell patch-clamp recordings under visual control. Patch pipettes with tip resistance of 4-6 MΩ were fabricated from capillary glass (1.50 mm outer diameter, 0.86 mm inner diameter) and filled with solution containing (in mM): 140 potassium gluconate, 5 NaCl, 0.1 EGTA, 10 HEPES, 2 Mg-ATP, and 0.3 Na_3_-GTP. Alexa 568 hydrazide dye was added to the patch-pipette solution (50 μM, Invitrogen, Carlsbad, CA, USA) as a color contrast to EGFP following whole-cell dialysis. Membrane potential was amplified (100x) and low-pass filtered (1 kHz) using a patch-clamp amplifier (EPC10, HEKA Elektronic, Holliston, MA, USA) and digitally acquired at 4 kHz (PowerLab 4/30, AD Instruments, Colorado Springs, CO, USA).

### Virus injection and fiber optic implantation

We anesthetized adult Dbx1;ArchT and Dbx1;CatCh (aged 8-20 weeks) mice via intraperitoneal injection of ketamine (100 mg/kg) and xylazine (10 mg/kg) and performed aseptic surgeries in the prone position using a stereotaxic frame. After exposing the skull, we performed either one (Dbx1;CatCh mice) or two (Dbx1;ArchT mice) 0.5-mm-diameter craniotomies in the range 6.95 to 7.07 mm posterior to bregma and 1.1 to 1.3 mm lateral to the midline suture.

In Dbx1;CatCh mice, we unilaterally injected an adeno-associated virus (AAV) immediately prior to fiber optic implantation to induce Flp-mediated recombination. We loaded an ultrafine, microvolume syringe (Neuros series, Hamilton, Reno, NV) with 120 μl of AAV-eSyn-FLPo (titer 10^13^ vg/ml, Vector Biolabs, Malvern, PA, USA). The syringe was lowered at 10 μm/s through the cerebellum and the virus was injected at the target site at approximately 60 nl/min. The syringe remained in place for 10 min before being retracted at 10 μm/s.

Both Dbx1;ArchT and Dbx1;CatCh mice were equipped with fiber optic appliances constructed by joining 1.27-mm-diameter ceramic ferrules (Precision Fiber Products, Milptas, CA, USA) with 105-μm-diameter 0.22 numerical aperture (NA) multimode fibers (Thorlabs, Newton, NJ, USA). We implanted fiber optic appliances bilaterally in Dbx1;ArchT mice and unilaterally in Dbx1;CatCh mice at a depth of 5.5 to 5.9 mm from bregma, which were secured with a cyanoacrylate adhesive (Loctite 3092, Henkel Corp., Rocky Hill, CT, USA). Dbx1;ArchT animals recovered for a minimum of 10 days before any further experimentation. Dbx1;CatCh mice recovered for a minimum of 21 days before further experimentation.

### Breathing measurements

After anesthetizing mice using 2% isoflurane we connected the ferrules of Dbx1;ArchT mice to a 589-nm laser (Dragon Lasers, Changchun, China). The ferrule of Dbx1;CatCh mice was connected to a 473-nm laser (Dragon Lasers). Mice recovered from isofluorane anesthesia for ~1 hr, and then we measured breathing behavior using a whole body plethysmograph (Emka Technologies, Falls Church, VA, USA) that allowed for fiberoptic illumination in a sealed chamber.

In a separate session, these same mice were lightly sedated via intraperitoneal ketamine injections (15 mg/kg minimum dose), which we titrated as needed to reduce limb movements but retain toe-pinch and blink reflexes. The maximum aggregate dose was limited to 50 mg/kg. Mice were fitted with a modified anesthesia mask (Kent Scientific, Torrington, CT, USA) to measure breathing.

We applied a circuit of positive pressure, with balanced vacuum, to continuously flush the plethysmograph with breathing air. The plethysmograph and the mask were connected to a 1-liter respiratory flow head and differential pressure transducer that measured airflow; positive airflow reflects inspiration in all cases. Analog breathing signals were digitized at 1 kHz (PowerLab).

### Optogenetic protocols

We applied 5 s bouts of light (either 473 or 589 nm) to Dbx1;ArchT and Dbx1;CatCh mice at graded intensities of 6.8, 8.6, and 10.2 mW. All ferrules were tested with a power meter prior to implantation to verify that illumination intensity did not vary more than 0.1 mW from the specified values. Bouts of light application were separated by a minimum interval of 30 s. We also applied 100 ms light pulses at a fixed intensity of 10.2 mW. We exposed each mouse to 85-200 pulses spaced at random intervals of between 1 and 5 s.

We applied 2 s bouts of 589-nm light (at the same intensities listed above) to rhythmically active slices. The fiberoptics were targeted to selectively illuminate the preBötC bilaterally but not the adjacent reticular formation.

### Data analyses

The airflow signal was band-pass filtered (0.1-20 Hz) and analyzed using LabChart 8 software (AD Instruments), which computes airflow (units of ml/s), respiratory rate (i.e., frequency, *f*, units of Hz), tidal volume (V_T_, units of ml), inspiratory time (T_i_), and minute ventilation (MV, units of ml/min). We computed statistics using GraphPad Prism 6 (La Jolla, CA, USA) and R: The Project for Statistical Computing (R, The R Foundation, Vienna, Austria) and prepared figures using Adobe Illustrator (Adobe Systems Inc., San Jose, CA, USA), GraphPad Prism 6, and IGOR Pro 6 (Wavemetrics, Lake Oswego, OR, USA). We analyzed the experiments in which 5 s light pulses were applied to the preBötC using paired t-tests, specifically comparing mean *f*, V_T_, and MV for control and illumination conditions at three different light intensity levels (i.e., at each laser strength tested, the pre-illumination ventilation serves as its own control).

We analyzed phase-response relationships of the breathing cycles perturbed by 100 ms-duration light pulses (see Figure 3C inset). The expected cycle period was measured from the unperturbed cycle immediately before the light pulse, which was defined as spanning 0-360° (Φ_Expected_). Cycle times were measured from the start of inspiration in one breath to the start of inspiration of the subsequent breath. For perturbed cycles, 100-ms light pulses were applied at random time points spanning the inspiration and expiration to test for phase shifts. Φ_Stim_ marks the phase at which the light pulse occurred. The induced cycle period (Φ_Induced_) was measured from the perturbed cycle. The perturbation of breathing phase, Φ_Shift_, was defined as the difference between Φ_Induced_ and Φ_Expected_. We calculated change in V_T_ and T_i_ in the perturbed breath compared to the expected breath normalized to the expected breath (refered to as, ΔV_T_ and ΔT_i_, respectively). Further, we calculated the phase shift of the breath following the perturbed breath (i.e., the cycle after Φ_Induced_) also with respect to Φ_Expected_; we refer to the phase of the subsequent breath Φ_N+1_. Measurements of Φ_Shift_, ΔV_T_, ΔT_i_, and Φ_N+1_ are all linked to a particular Φ_Stim_ within the interval 0-360°. To analyze group data we sorted Φ_Stim_ into 12 equally sized 30° bins. We computed the mean and standard deviation (SD) for Φ_Shift_, ΔV_T_, ΔT_i_, and Φ_N+1_ within each bin, which we then plotted in phase-response curves along with values calculated from wild type littermates. A Tukey’s HSD to test was used to evaluate how unlikely it would have been to obtain mean Φ_Shift_, ΔV_T_, ΔT_i_, and Φ_N+1_ for each bin if the optogenetic perturbations had commensurate effects on Dbx1;ArchT (or Dbx1;CatCh) mice and wild type littermates.

### Histology

After experimentation we verified in all animals that fiber optic tips were within 500 μm of the dorsal preBötC border, which could be identified via well-established anatomical criteria in combination with either ArchT-EGFP or CatCh-EYFP fusion protein expression in reporter mice (Figure 1C). We administered a lethal dose of pentobarbital (100 mg/kg i.p.) and then transcardially perfused the mice with 1x PBS followed by 4% PFA in PBS. The neuraxes were removed and post-fixed overnight in 4% PFA, and later sliced in 50-μm contiguous transverse sections using a vibratome. Free-floating sections were stained using NeuroTrace 530/615 red fluorescent nissl stain (Invitrogen) for 1 hr, rinsed in PBSand then cover-slipped using Vectashield (Vector Labs, Burlingame, CA, USA). Tissue sections were visualized using bright-field and confocal microscopy. Images were arranged as mosaics and brightness and contrast were adjusted uniformly across the entire ensemble image using the public domain software package ImageJ. Images were not manipulated in any other way.

## Acknowledgements

National Institutes of Health grant R01 HL104127 (PI: Del Negro) supported this work.

---

**Figure 1 - figure supplement 1.**
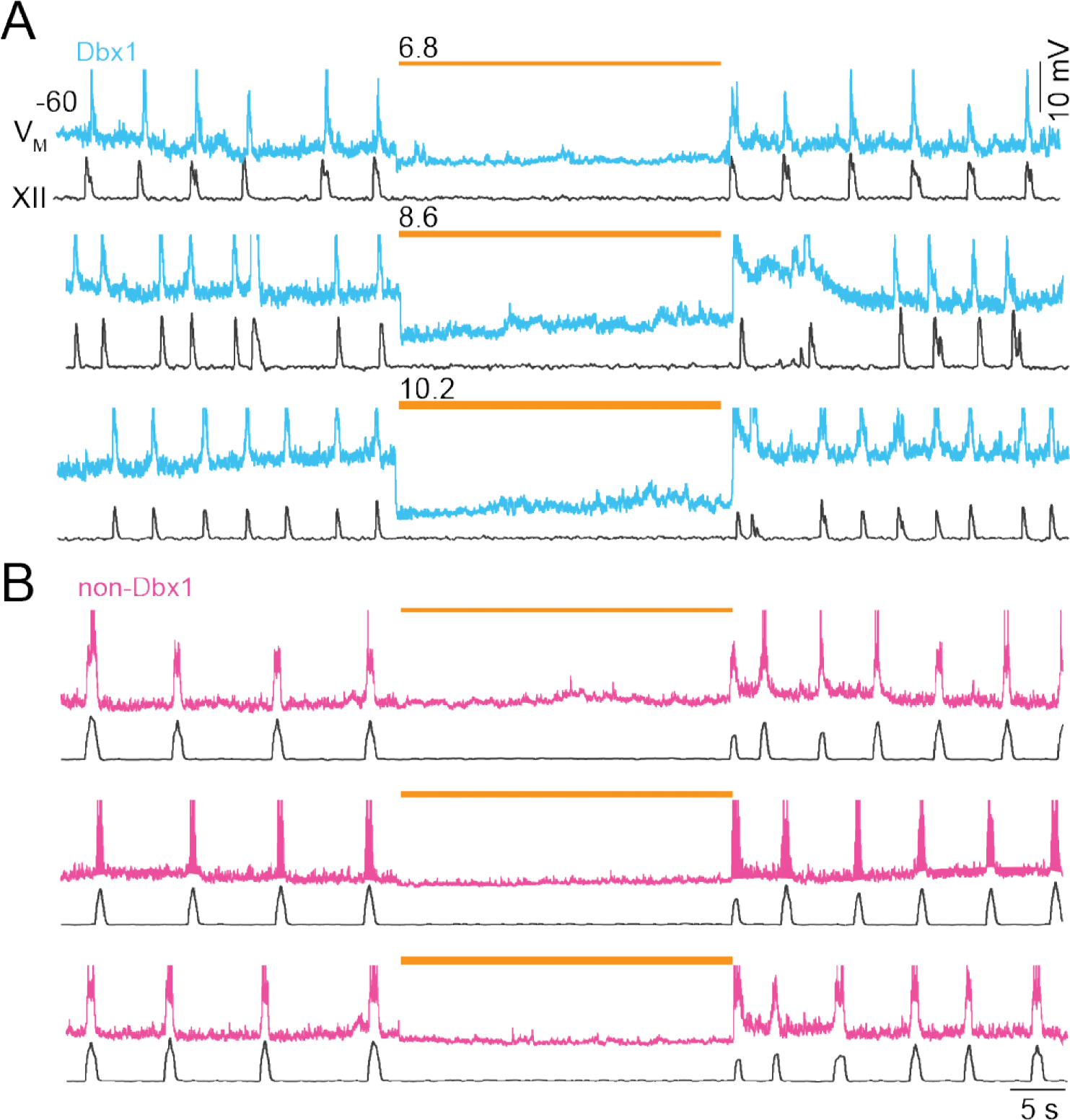
Photoinhibition of preBötC neurons *in vitro*. **A**, Membrane trajectory of an ArchT-expressing Dbx1 preBötC neuron (V_M_, cyan traces) in a rhythmically active slice preparation with inspiratory motor output recorded from the XII nerve rootlet. **B**, Membrane trajectory of a non-Dbx1, non-ArchT-expressing preBötC neuron (V_M_, magenta traces) with XII motor output. Light pulses (30 s) were applied bilaterally to the preBötC at three intensities (units of mW) in A and B. Yellow line thickness corresponds to light intensity, which is also annotated above each line. Voltage and time calibrations apply to A and B, including baseline membrane potential of −60 mV. Action potentials have been truncated for display to emphasize the trajectory around the baseline membrane potential.

**Figure 2 - figure supplement 1.**
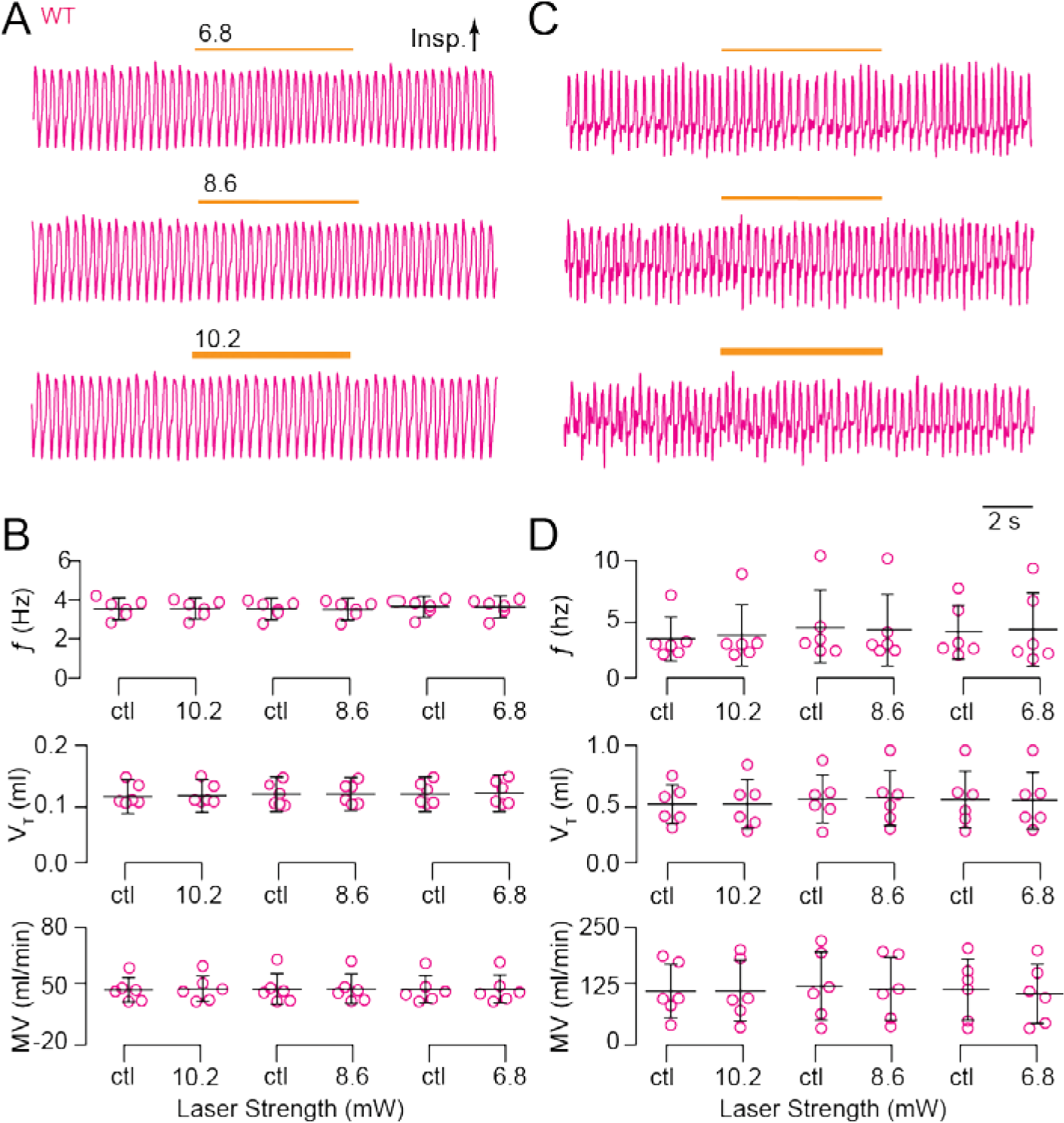
Light application to the preBötC does not affect breathing in wild type Dbx1;ArchT littermates. **A**, Airflow traces from a sedated mouse exposed to 5-s bouts of bilateral preBötC illumination at three intensities (units of mW). Yellow line thickness corresponds to light intensity, which is also annotated above each line. **B**, Group data from experiments in A quantifying *f*, V_T_ and MV in response to light application. Symbols show mean *f*, V_T_, and MV in each mouse. Bars show the mean and SD for all animals tested (n = 6). Control measurements are labeled ‘ctl’; numerals indicate light intensity. **C**, Airflow traces from an awake unrestrained mouse exposed to 5-s bouts of unilateral preBötC illumination at three intensities (units of mW). Yellow line thickness corresponds to light intensity; annotations mach those in A. **D**, Group data from experiments in C quantifying *f*, V_T_ and MV in response to light application. Symbols show mean *f*, V_T_, and MV in each mouse. Bars show the mean and SD for all animals tested (n = 6). Control measurements are labeled ‘ctl’; numerals indicate light intensity.

**Figure 3 - figure supplement 1.**
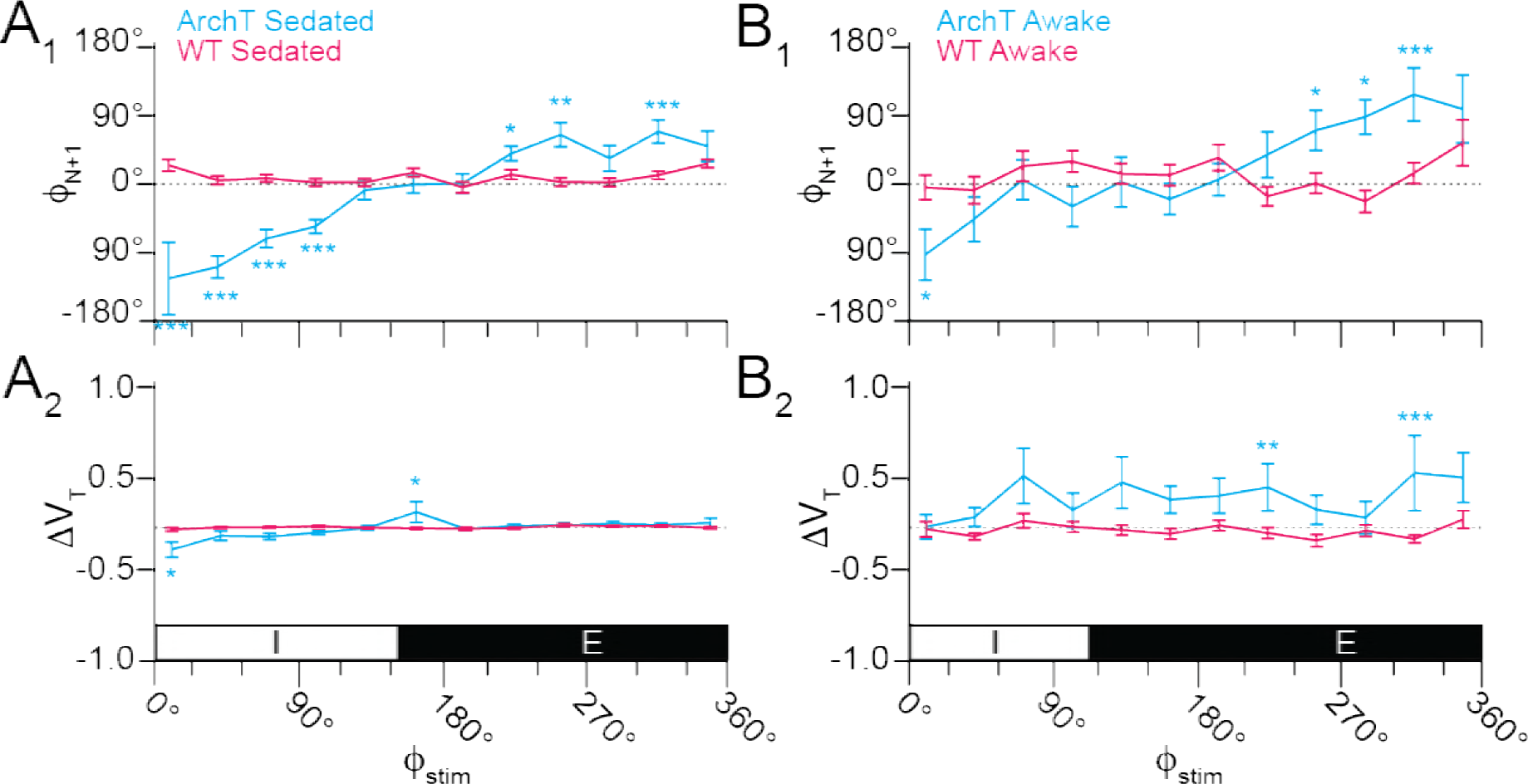
Effects of brief photoinhibition on V_T_ and Φ_N+1_ in Dbx1;ArchT mice (n = 5 in A, n = 6 in B, cyan) and wild type littermates (n = 6, magenta). **A_1_**, Phase-response curve plotting Φ_N+1_ vs. Φ_Stim_ throughout the breathing cycle in sedated mice. **A_2_**, Phase-response curve for changes in V_T_ following brief photoinhibition (i.e., the perturbed breath) in the same cohort of sedated mice. The abscissa marks the inspiratory (I, 0-150°) and expiratory (E, 150-360°) phases of the breathing cycle (0-360°), which applies to A_1_ and A_2_. **B_1_**, Phase-response curve plotting Φ_N+1_ vs. Φ_Stim_ in awake unrestrained mice. **B_2_**, Phase-response curve for ΔV_T_ vs. Φ_Stim_ in the same cohort of awake unrestrained mice. The abscissa marks the inspiratory (I, 0-110°) and expiratory (E, 110-360°) phases of the complete breathing cycle (0-360°), which applies to B_1_ and B_2_.

**Figure 4 - figure supplement 1.**
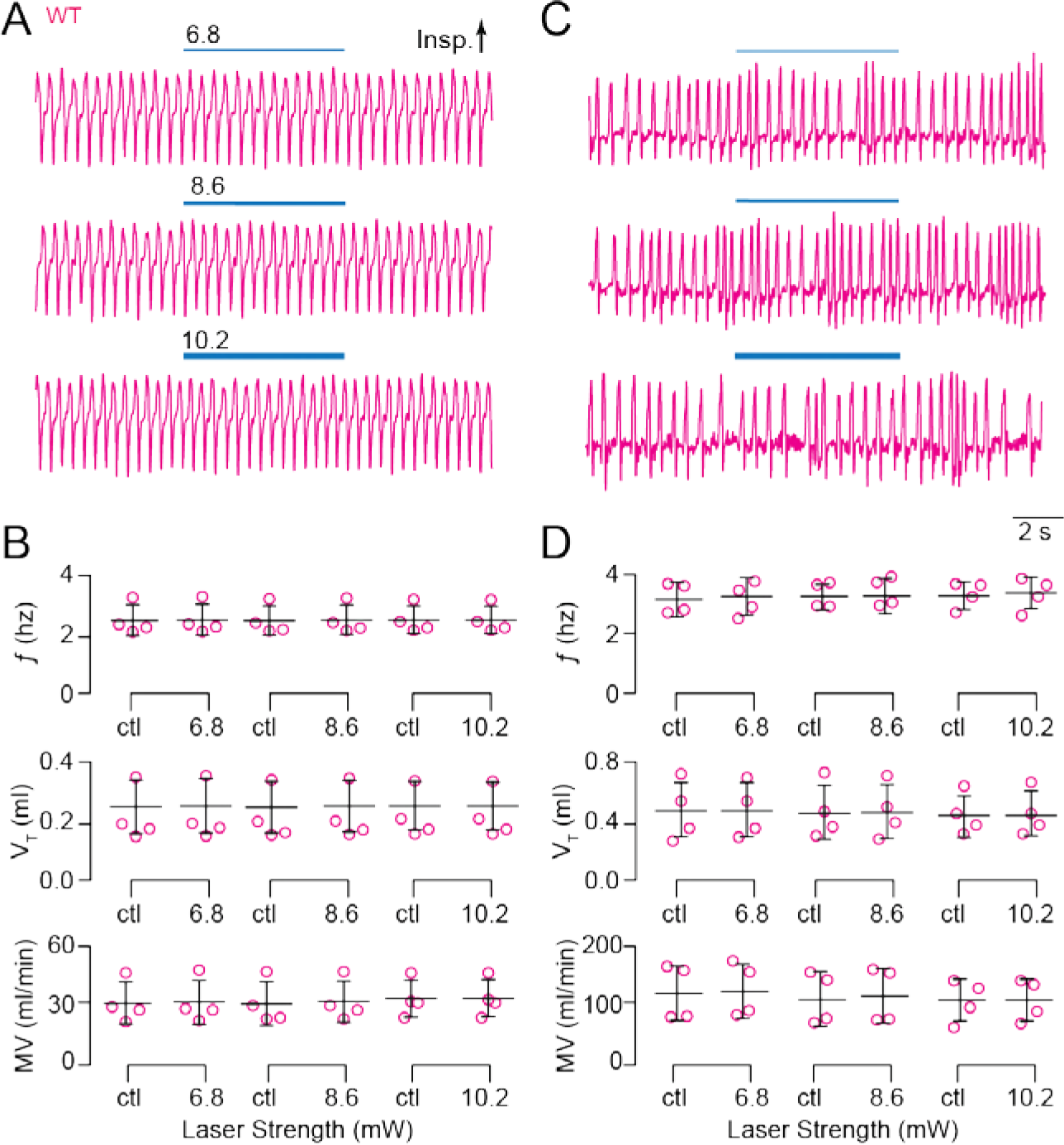
Light application to the preBötC does not affect breathing in wild type Dbx1;CatCh littermates. **A**, Airflow traces from a sedated mouse exposed to 5-s bouts of unilateral preBötC illumination at three intensities (units of mW). Cyan line thickness corresponds to light intensity, which is also annotated above each line. **B**, Group data from experiments in A quantifying *f*, V_T_ and MV in response to light application. Symbols show mean *f*, V_T_, and MV in each mouse. Bars show the mean and SD for all animals tested (n = 4). Control measurements are labeled ‘ctl’. **C**, Traces from an awake unrestrained mouse exposed to 5-s bouts of unilateral preBötC illumination at three intensities. Cyan line thickness corresponds to light intensity; annotations mach those in A. **D**, Group data from experiments in C quantifying *f*, V_T_ and MV in response to light application. Symbols show mean *f*, V_T_, and MV in each mouse. Bars show the mean and SD for all animals tested (n = 6). Control measurements are labeled ‘ctl’; numerals indicate light intensity.

**Figure 5 - figure supplement 1.**
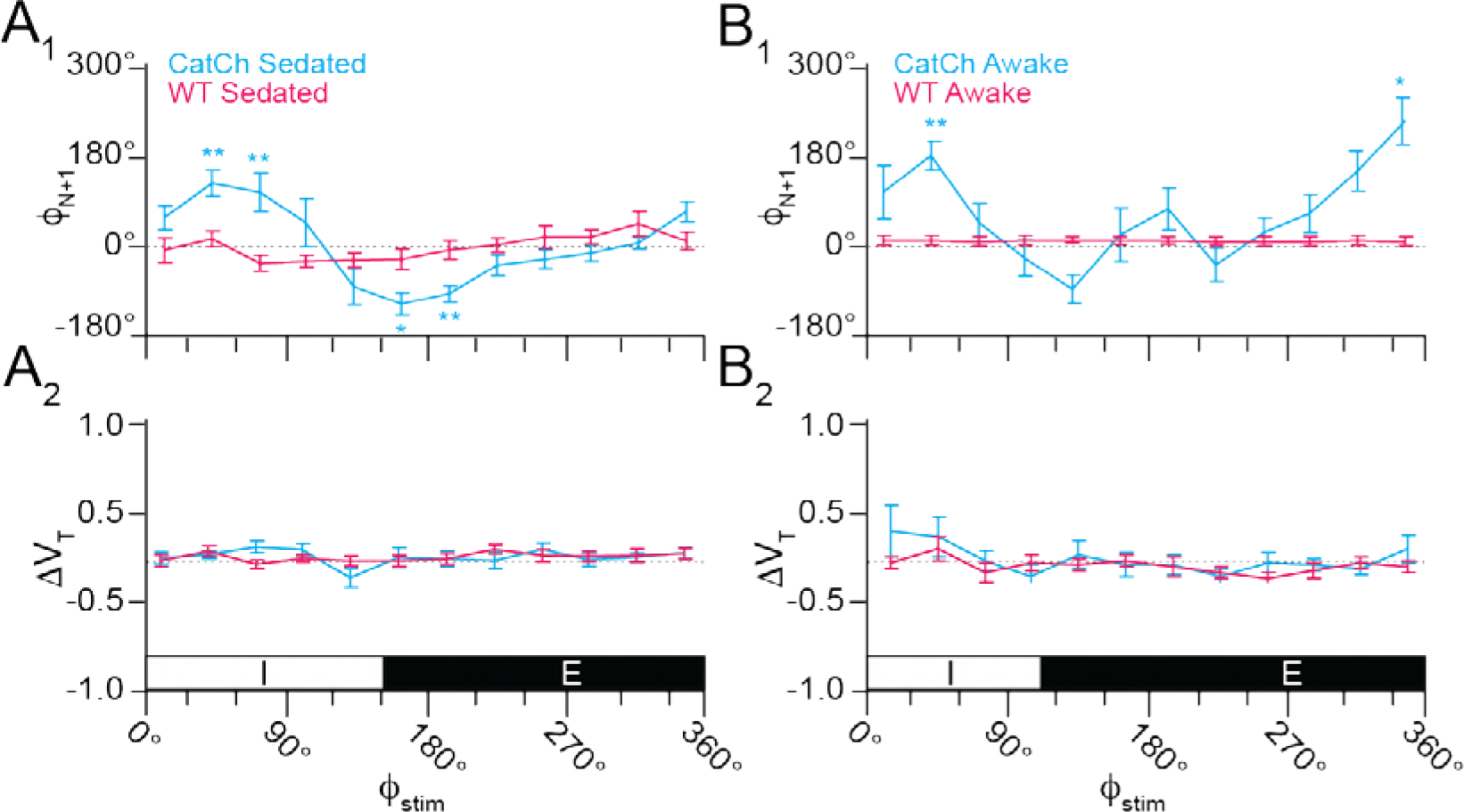
Effects of brief photostimulation on V_T_ and Φ_N+1_ in Dbx1;CatCh mice (n = 4, cyan) or wild type littermates (n = 4, magenta). **A_1_**, Phase-response curve plotting Φ_N+1_ vs. Φ_Stim_ throughout the breathing cycle in sedated mice. **A_2_**, Phase-response curve for changes in V_T_ following photostimulation (i.e., the perturbed breath) in the same cohort of sedated mice (n = 4). The abscissa marks the inspiratory (I, 0-150°) and expiratory (E, 150-360°) phases of the breathing cycle (0-360°), which applies to A_1_ and A_2_. **B_1_**, Phase-response curve plotting Φ_N+1_ vs. Φ_Stim_ in awake unrestrained mice. **B_2_**, Phase-response curve for ΔV_T_ vs. Φ_Stim_ in the same cohort of awake unrestrained mice. The abscissa marks the inspiratory (I, 0-110°) and expiratory (E, 110-360°) phases of the complete breathing cycle (0-360°), which applies to B_1_ and B_2_.

